# Neuroinflammation increases oxygen extraction in a mouse model of Alzheimer’s disease

**DOI:** 10.1101/2023.10.16.562353

**Authors:** Chang Liu, Alfredo Cardenas-Rivera, Shayna Teitelbaum, Austin Birmingham, Mohammed Alfadhel, Mohammad A. Yaseen

## Abstract

Neuroinflammation, impaired metabolism, and hypoperfusion are fundamental pathological hallmarks of early Alzheimer’s disease (AD). Numerous studies have asserted a close association between neuroinflammation and disrupted cerebral energetics. During AD progression and other neurodegenerative disorders, a persistent state of chronic neuroinflammation reportedly exacerbates cytotoxicity and potentiates neuronal death. Here, we assessed the impact of a neuroinflammatory challenge on metabolic demand and microvascular hemodynamics in the somatosensory cortex of an AD mouse model. We utilized in vivo 2-photon microscopy and the phosphorescent oxygen sensor Oxyphor 2P to measure partial pressure of oxygen (pO2) and capillary red blood cell flux in cortical microvessels of awake mice. Intravascular pO2 and capillary RBC flux measurements were performed in 8-month-old APPswe/PS1dE9 mice and wildtype littermates on days 0, 7, and 14 of a 14-day period of lipopolysaccaride-induced neuroinflammation. Before the induced inflammatory challenge, AD mice demonstrated reduced metabolic demand but similar capillary red blood cell flux as their wild type counterparts. Neuroinflammation provoked significant reductions in cerebral intravascular oxygen levels and elevated oxygen extraction in both animal groups, without significantly altering red blood cell flux in capillaries. This study provides evidence that neuroinflammation alters cerebral oxygen demand at the early stages of AD without substantially altering vascular oxygen supply. The results will guide our understanding of neuroinflammation’s influence on neuroimaging biomarkers for early AD diagnosis.

## Background

Neuroinflammation is a key contributor to the pathogenesis of Alzheimer’s disease (AD) and other neurodegenerative disorders (1–4). Activated microglia and astrocytes constitute the principal mediators of neuroinflammation through overproduction of proinflammatory cytokines, chemokines, and reactive oxygen species (ROS), while vascular endothelial cells contribute through expression of vascular and cellular adhesion molecules (5–9). In addition to their vital roles in neuroinflammation, these cells are also crucial components of the neuro-glio-vascular unit (NGVU), which act in concert to locally regulate microvascular blood flow to satisfy the brain’s dynamic, heterogeneous energy demands (10). Disruptions to the NGVU account for many of AD’s other preclinical hallmarks, including reductions to cerebral perfusion, impaired glucose and oxygen metabolism, and neurovascular decoupling (11–15).

A growing body of evidence highlights the causative and exacerbating roles of inflammation in the preclinical etiology of AD and other neurodegenerative disorders (2,3,16–20). In diseases such as multiple sclerosis and Parkinson’s disease, a single immune stimulus can initiate a chronic neuroinflammatory phenotype in microglia and promote self-perpetuating neurotoxicity (20,21). In the earliest stages of AD, microglial activation reportedly precedes formation of Amyloid β plaques (22,23). Proinflammatory cytokines and chemokines promote synaptic loss, Aβ aggregation, increased oxidative stress, and mitochondrial dysfunction (20,24). Because it potentiates cerebrovascular pathology, mitochondrial dysfunction, and Aβ accumulation, the critical role of neuroinflammation in preclinical AD’s destructive positive-feedback cascade continues to gain recognition (13,20). Consequently, the influence of neuroinflammation on cerebral blood flow and energy metabolism warrants particular consideration (25–27). In peripheral tissues, immunological studies have demonstrated that systemic inflammatory responses alter oxygen and glucose metabolism, mitochondrial membrane permeability and oxidative phosphorylation, and adenosine triphosphate (ATP) consumption (28,29,19). Similarly, in the central nervous system, neuroinflammation stimulates production of ROS while inducing a glycolytic shift and mitochondrial fission or fusion in microglia and astrocytes, all of which significantly alters oxygen and glucose consumption and associated aerobic ATP production (28,30–33). Alterations to morphology, function, and metabolism of astrocytes, vascular endothelial cells, and microglia likely confound their roles in the NGVU for moderating supply of oxygen and other metabolites. The corresponding surge of neutrophils, monocytes, and lymphocytes and their associated cytokines in the bloodstream can also disrupt circulation through microvascular networks (34,35). Consequently, neuroinflammation directly impacts translatable biomarkers of AD progression through by altering the dynamic cerebral blood flow and oxygenation signals that serve as the basis for functional MRI, near-infrared spectroscopy, and positron emission tomography (36,37). Previous neuroimaging studies reported empirical correlations between increased neuroinflammation and reduced brain energy metabolism in AD patients and in preclinical AD mouse models (26,27), as well as studies of functional somatic syndrome and mild traumatic brain injury (38,39). To precisely understand how it can aggravate and accelerate clinically observable signatures of AD’s pathogenesis, the impact of neuroinflammation’s pronounced influence on cerebral energetics and hemodynamics requires investigation at the microscopic scale in live brains.

Neuroinflammatory processes reportedly exert both beneficial and harmful effects during disease progression (1,40). These double-edged effects have motivated several studies exploring the potential effects of enhancing or repressing a pro-inflammatory response (41). In this study, we investigated whether changes in cortical oxygenation, microvascular hemodynamics, and neuroinflammation are evident at the early stages of AD progression in an AD mouse model. We also explored the presumptive catalytic role of a pronounced neuroinflammatory stimulus on neurovascular and metabolic dysregulation in AD progression. We applied in vivo two-photon phosphorescence lifetime imaging microscopy (2P-PLIM) with the novel oxygen-sensitive sensor, Oxyphor 2P, to measure oxygen partial pressure (pO2) and capillary red blood cell flux (RBC flux) in awake APPswe/PS1dE9 mice and wildtype controls at age 7-8 months (42–44). The 2P-PLIM method uniquely enables nondisruptive, longitudinal characterization of pO2 and cerebral hemodynamics within individual microvessels. Previously, two-photon microscopy has revealed widely-varying neuronal and microvascular densities widely across different cortical layers, supporting observed laminar variations of cortical energy metabolism (45–50). Capitalizing on the technique’s penetration depth and unparalleled spatial resolution, we quantified layer-specific differences in vascular oxygen extraction and capillary flux provoked by a prolonged inflammatory threat. Microvascular pO2 and capillary RBC flux measurements were collected from cortical layers I-IV before, during, and after 14 days pro-inflammatory stimulus, induced by daily intraperitoneal injection of the bacterial endotoxin Lipopolysaccharide (LPS) (51–55). The measurements were used to compute layer-specific changes in oxygen extraction and capillary red blood cell flux. Before induced inflammation, AD mice show lower amounts of oxygen extraction compared to WT mice, reflecting reduced metabolic demand in AD brain tissue. LPS-induced inflammation activated provoked increased cytokine expression from microglia and astrocytes and substantially increased oxygen extraction at all cortical depths in both AD and control groups without altering RBC flux dynamics. Our results show that cerebral oxygen consumption is affected more significantly than vascular oxygen supply at early stages of AD, consistent with prior studies. A pro inflammatory stimulus increases oxygen extraction in the brain tissue without significantly altering vascular oxygen supply in the early stages of preclinical AD. The study emphasizes the interdependent nature of metabolic and neuroimmune alterations that transpire during preclinical AD.

## Methods

### Experimental design

All experiments were performed in accordance with ARRIVE guidelines for animal care, under a protocol approved by the Northeastern University Institutional Animal Care and Use Committee. Female APP/PS1dE9 mice and their age-matched wild-type littermates are used in this study (*n*=10 each, The Jackson Laboratory, MMRRC Strain # 034829-JAX). The double transgenic AD mouse co-expresses the Swedish mutation of amyloid precursor protein (APP) and a mutant human presenilin 1 gene (PS1-dE9) and accumulates beta-amyloid deposits in the cortex at 6 months (56). Mice were prepared with cranial window surgery at 6 months of age followed by habituation training of head-fixed restraint. Prior to LPS-induced systemic inflammation, “baseline” cerebral intravascular oxygenation and hemodynamics were measured in AD (8.19 months old on average) and WT (7.95 months old on average) mice. To induce systemic inflammation, 0.25 to 1 mg/kg, 1 mg/ml PBS-LPS solution (LPS from Escherichia coli O55:B5, Product Number: L4005, CAS-No. 93572-42-0, Sigma Aldrich) was administered to all mice via intraperitoneal injection daily for two weeks. Animals’ body weights and sickness behavior (i.e., posture, activity) were monitored daily to assess the sickness of the animal as well as the efficacy of the LPS. Two AD and three WT mice had gasping and low body temperature after the first two to three injections of LPS and did not survive LPS administration, and thus were excluded from the analyses. Microvascular pO2 and capillary RBC flux were measured after one week and two weeks of LPS injection. Following these measurements, mice were euthanized and brains (*n*=3 in each cohort) were harvested for immunohistochemical analysis.

### Animal preparation

Following a procedure described previously (57), mice underwent cranial window surgery at age 6 months. Succinctly, ∼4 hours before the surgery, the mouse was given IP injection of dexamethasone (4.8 mg/kg at 4 mg/ml) and cefazolin (0.5 g/kg at 200 mg/ml) to decrease brain inflammation and edema during and after surgery. During the surgery, the mouse was anesthetized with isoflurane (3∼4% for induction and 1∼2% for maintenance). Mouse head was secured on a stereotaxic instrument (David Kopf Instruments, CA, USA), and a heating pad (Harvard Apparatus, MA, USA) was placed under the mouse body to maintain physiological temperature. Hair and skin were removed to expose the skull. Using a dental drill, a 3-mm-diameter cranial window was made over the left somatosensory cortex (ML: -3mm, AP: -2 mm to bregma). The skull was removed and the dura was left intact. A custom-made glass coverslip plug (OD=5mm, ID=3mm) was placed on top of the exposed dura and was sealed with superglue (57). A customized head post was glued to the right hemisphere with Loctite 401. The skull was sealed with dental acrylic. To prevent dehydration, 100 µl, 5% dextrose sodium solution was injected subcutaneously. The mouse was singly housed and supported with antibiotics (40/8 mg/ml, sulfametoxazole/trimethiporim (SMX-TMP) and 50 mg/ml, carprofen in drinking water) continuously for five days for post-operative recovery and to minimize surgical infection. Buprenorphine (0.05 mg/kg at 0.03 mg/ml) was administered subcutaneously as analgesic. Headpost restraint training started at 7 days post-surgery using a custom-made imaging cradle. During training, animals were conditioned to tolerate head immobilization for durations lasting from 5 mins to 2 hours to get the animal acclimated to the two-photon imaging experiments. Milk was given as a reward during the training. Imaging took place one month after the surgery to ensure full recovery from surgery-induced inflammatory response (58,59).

### Cerebral intravascular pO2 measurements

Approximately 30 minutes before imaging experiments, the oxygen-sensitive phosphorescent probe, Oxyphor 2P (43), was administered to mice via retro-orbital injection (80 µM, 50 - 60 µl) under anesthesia (∼ 2% isoflurane). The probe uniformly distributes inside the vasculature and labels blood plasma, allowing 2-3 hours’ intravascular imaging. The probe has a maximum two-photon absorption at λ=950 nm and emits red-shifted photons with a central wavelength of 758 nm (43). After recovering from anesthesia, the mouse was positioned in a customized cradle with its head fixed and cranial window surface leveled. Condensed milk was provided every 30 minutes.

Phosphorescence lifetime measurements were performed using a commercial two-photon microscope (Ultima2P plus, BrukerNano, Inc.), a tunable ultrafast pulsed laser (InSightX3+, Spectral-Physics, US) with a repetition rate of 80 MHz was tuned at 950 nm, and a water-immersion 25X objective (N25X-APO-MP - 25X Nikon, 1.10 NA, 2.0 mm WD) (Fig. 1A). The objective was heated to ∼36.6 °C with an objective heater and temperature controller (ALA Scientific Instruments, Inc.) to prevent heat loss from the cranial window and to eliminate temperature-related variability in phosphorescence lifetime (60,43). For each animal, intravascular pO2 within a 500 μm field of view (Fig. 1B, red box) was measured at multiple depths in the cortex. At the beginning of the experiment, the FOV was raster-scanned for 7 to 15 seconds, and the photon emitted from each pixel after each pulsed laser was counted through time-correlated single-photon counting (TCSPC) (61). Emitted photons from each pixel were summed to generate a 2D image revealing the vasculature of the brain cortex (Fig. 1C). Intravascular locations were selected based on the 2D image for 2P-PLIM measurements and pO2 quantitation. Each pointwise PLIM measurement cycle consisted of a 10 µs-long excitation pulse, followed by a 290-µs-long collection of the emitted photons via time-correlated single photon counting (TCSPC). The 300-us cycle was repeated 2000 times (∼ 0.6s). The TCSPC hardware (SPC-150N, Becker & Hickl, GmbH), accumulated a time-binned histogram of emitted photons over the 290 μs decay period, yielding an exponential distribution of the emitted photons with respect to the starting time of the excitation pulse (61). Histograms were collected from the 2000 cycles to yield an exponential decay with high signal-to-noise ratio (SNR), allowing for accurate lifetime calculation via nonlinear curve fitting. The 2000 cycles were iterated 20 times in all selected measurement locations, such that 20 phosphorescence decays were acquired for each intravascular location. For each intravascular location, the 20 phosphorescence decays were averaged to get the mean phosphorescence decay before lifetime and pO2 calculation. Fig. 1D shows the pO2 color-coded mean phosphorescence decays measured inside capillary locations in Fig. 1c. In each mouse, pO2 in diving arteriole and venules, and capillaries were measured at subpial depths of z=50, 100, 200, 300, 400, 500 µm. Additional file 1: Supplementary Table 1 shows the number of arterioles and venules being measured for pO2 in both animal cohorts.

**Fig. 1.**
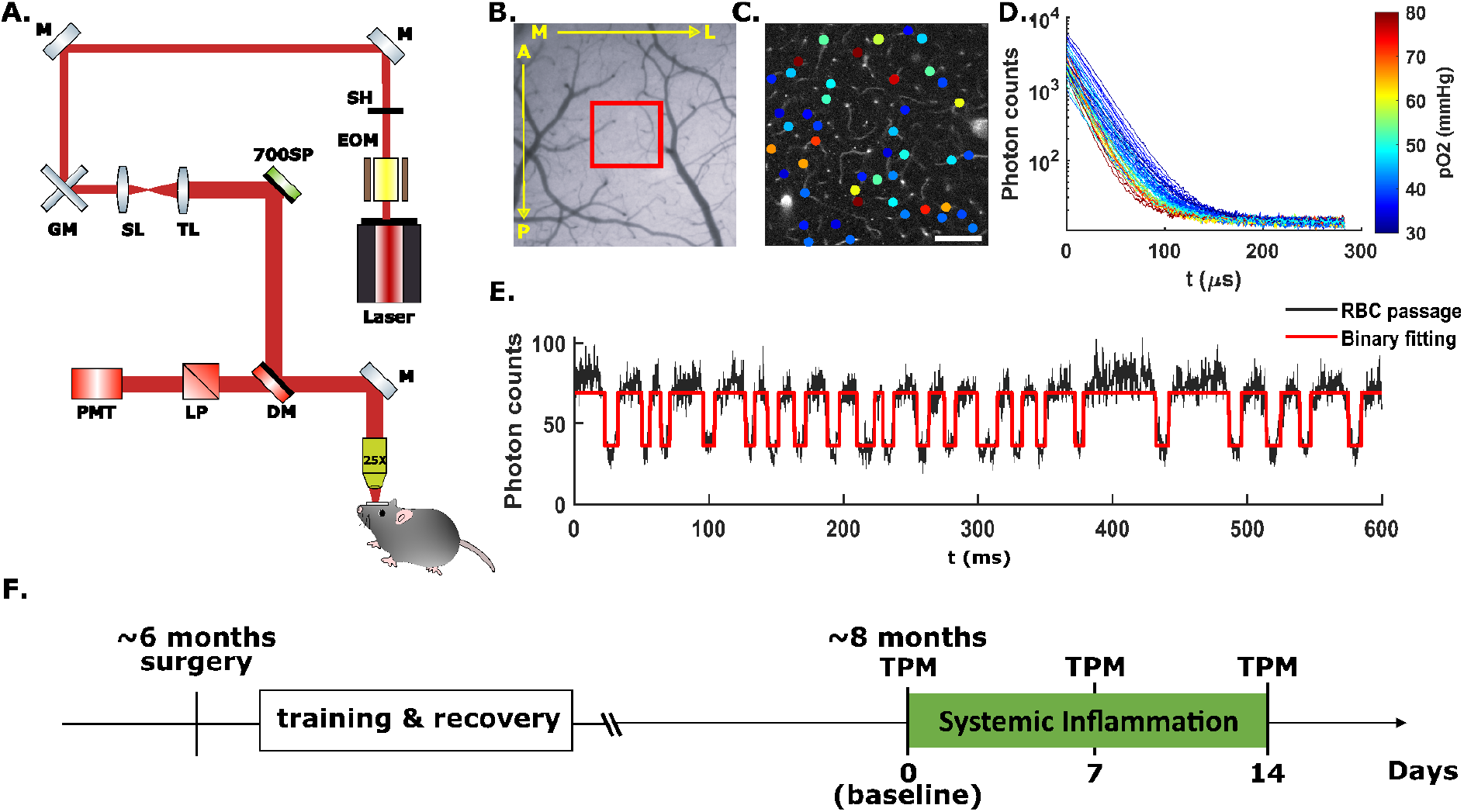
Experimental design and data acquisition. **(A)** Imaging set up for 2P-PLIM measurement in head-fixed, awake mice. 80 MHz, ultrafast pulsed laser tuned to 950 nm. The excitation pulsed laser is reflected by a 700 nm short pass filter and a 900 nm short pass filter and is focused to the imaging field through a 25X heated, water-immersive objective. The emission light is reflected and transmitted through the 900 SP filter and the 640 long-pass emission filter, and then detected by a photon-counting PMT. **(B)** Wide-field imaging of an AD mouse cranial window in the left somatosensory cortex. The red box indicates the region (Field of View: 474.27×474.27 µm^2^) being measured by two-photon microscopy. A: anterior, P: posterior, M: medial, L: lateral. **(C)** Two-photon survey scan image of the red-boxed field of view in **(B)** at cortical depth z=200 µm, scale bar = 100 µm. Colored dots are points of measurement of intravascular pO2 and RBC flux, color-coded by pO2 values. **(D)** Time-resolved phosphorescence emission of the corresponding pixels in **(C)** when the laser is off. Decays are color-coded by pO2 values. **(E)** Time-integrated phosphorescence intensity inside a single capillary over 600 ms (corresponding to 2000 PLIM measurements). Black curve is the time-integrated photon total counts. Local minima correspond to individual, non-labelled erythrocytes passing through the capillary (black). The timelapse intensity curve is thresholded (red) to count the number of RBCs (valleys of the intensity curve) over time. **(F)** Experimental timeline. EOM: electro-optic modulator. SH: shutter. M: mirror. GM: galvanometer scanner. SL: scan lens. TL: tube lens. SP: short-pass filter. LP: long-pass filter. DM: dichroic mirror. PMT: GaAsP photomultiplier tube.

### Cerebral capillary red blood cell flux measurements

Red blood cell flux was measured at the same capillary measurement locations using the time-binned phosphorescence photons of the 2P-PLIM pO2 measurements. Because RBCs do not get labeled by the Oxyphor2P probe, the measured phosphorescence signal decreases as individual RBCs pass through the focal point in capillaries. Consequently, the time-resolved phosphorescence intensity trace features “valleys” representing the passage of individual erythrocytes (62). With 2P-PLIM, phosphorescence emission photons of each 300 us cycle were binned to get the phosphorescence intensity. With 2000 cycles, a 0.6-s-long phosphorescence intensity time course I(t), (t=0 to 0.6 s, dt = 300 us) was obtained for each measured capillary (Fig. 1E). For each capillary segment, the mean RBC flux over all repetitions was calculated. Then the mean RBC flux of all measured capillary segments at all imaging depths were grouped into cortical layers, as described previously (45,44). The average RBC flux, standard deviation, and coefficient of variation (CV) in each cortical layer were calculated. The layer-specific results were averaged within animal cohorts to obtain the cortical layer-dependent mean RBC flux in AD and WT mice. For two WT mice, baseline RBC flux values were calculated using fluorescence measurements of dextran-conjugated FITC (70 kDa, Sigma Aldrich) on a separate day. Additional file 1: Supplementary Table 2 shows the number of capillaries measured for RBC flux for both animal cohort. Fig. 1F shows the experimental timeline.

### Calculation of pO2, SO2 and coefficient of variation

Intravascular pO2 was calculated in each measurement location from the computed lifetimes of the measured phosphorescence decays. For each measured location, the mean phosphorescence decay was calculated by averaging over the 20 repetitions. In accordance with the established analysis protocol for Oxyphor 2P (43), the first 5 µs of the recorded PLIM decays were discarded. The remaining 285 µs of the averaged phosphorescence decay was fitted using a mono-exponential decay function 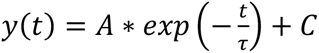, and the decay rate *τ* represents the phosphorescent lifetime. pO2 was calculated from τ using a calibration equation similar to the Stern-Vollmer relation (43). In some experiments, abnormal phosphorescence decays occur due to animal motions. To exclude these instances, the total photon of each phosphorescence decay was calculated and the decay curves with outlying total photons were rejected using Grubb’s test (63) (MATLAB) before calculating the mean phosphorescence decay of the measured point. In each mouse, intravascular pO2s were grouped into cortical layers based on subsurface measurement depth. At each cortical layer, mean pO2 and the coefficient of variation (CV) were calculated. Layer-dependent pO2s and CVs were then averaged over all measured animal samples in each group (AD & WT) to obtain the mean pO2s and CVs of AD and WT animals. pO2 was converted into oxygen saturation of hemoglobin (SO2) using the Hill equation with coefficients for C57BL/6 mice (h=2.59, P50 = 40.2 mmHg) (64). Layer-dependent SO2 for AD and WT were calculated using the same steps described in the calculation of layer-dependent pO2.

### Calculation of cortical depth-dependent oxygen extraction (DOE)

The amount of oxygen extracted from the cerebral capillary bed was estimated using the cortical depth-dependent oxygen extraction (DOE, %). In each cortical layer, the mean arteriole SO2 and venule SO2 was calculated through averaging over all measured arteriole and venule SO2s, and the oxygen extraction of the corresponding layer was calculated as the difference between the arteriole SaO2 and venule SvO2. DOE over all animals were averaged.

Cerebral oxygen extraction fraction (OEF) reflects the intricate balance between brain tissue’s metabolic demand and metabolite supply from the cerebral vasculature. This metric is used extensively in clinical neuroimaging studies as a biomarker for cognitive impairment. OEF is generally calculated as the arteriovenous difference in SO2, normalized by the arterial SO2. Typically, the arterial SO2 is considered constant at ∼98%, indicating complete saturation of O2 to RBCs. Because we observed significant reductions in arterial SO2 after inflammation, we opt to present our data as depth-dependent oxygen extraction, which indicates the non-normalized AV difference in SO2.

### Vessel Identification

During imaging experiments, large pial vessels were traced along the penetrating path from the pial surface to the deepest visible imaging plane (around 400 µm to 550 µm) for arteriole and venule pO2 measurements. Arterioles and venules were distinguished later using the pO2 values. According to the values reported in awake and anesthetized animals (46,65), we classified vessels with surface pO2 (measured at 50 *µm* below the pial surface) higher than 66 mmHg as arterioles, and those below 66 mmHg as venules. Note that the arterioles and venules were determined using the baseline pO2 values (pre LPS-induced inflammation). Capillaries were identified based on morphology and tortuosity. In each imaging plane, all visible capillary segments were selected for pO2 and RBC flux measurement. Primary branches from the penetrating vessels were considered precapillary arterioles or postcapillary venules and were excluded from the capillary pO2 calculation during data analysis.

### Tissue collection and immunofluorescence staining

After the last imaging session, a subgroup of AD and WT mice (*n*=3 in each cohort) were intracardially perfused with heparin-dissolved 1X PBS followed by 4% paraformaldehyde (PFA). Brains were collected and stored in 4% PFA at 4 °C. We also collected the brains of age-matched AD and WT mouse that never received LPS as the control group (*n*=1 in each cohort). The cerebrums were sectioned into 3 to 4 coronal slices (∼ 2 mm thickness) and stored in 75% ethanol. The sections were processed and stained with immunofluorescent antibodies by Dana-Farber/Harvard Cancer Center Specialized Histopathology Services (SHS) Core (Boston, MA.). Briefly, nuclei were stained with DAPI. Microglia and astrocytes were stained with Iba1-conjugated Alexa Fluor 594 and GFAP-conjugated Alexa Fluor 488, respectively. The stained brain slices were mounted on the glass slide for microscopic imaging.

### Confocal imaging and immunofluorescence analysis

Harvested, stained brain slices were imaged with a laser scanning confocal microscopy (Zeiss, LSM, 800) using a 10X objective. A 709 ×709 µm^2^ region of interest (ROI) around the somatosensory cortex in both hemispheres were selected. Two to six ROIs from 3 brain slices were imaged for each mouse. DAPI, Alexa Fluor 488, and Alexa 594 were excited with laser wavelength at 405 nm, 488 nm, and 561 nm. Pinhole sizes were set at 1 airy disk unit for each wavelength. The excitation power and detector gain were determined and fixed for all slices using one of the brightest slices to avoid intensity saturation. Z stack images were collected with a thickness of 40 µm and a step size of 3.96 µm. To quantify the expression of GFAP and Iba-1 in astrocytes and microglia cells, respectively, we conducted densitometry analysis. Maximum intensity projection (MIP) was calculated over the full 40 µm tissue thickness. The MIP images were binarized using an intensity threshold determined by the intensity histogram. Identical thresholding values for GFAP and Iba-1 were used among all MIP images for accurate calculation of pixel total intensity. Percentage area (PA) of GFAP and Iba-1 positive cells were calculated as the ratio between the number of pixels with the intensity above the threshold to the total number of pixels. For each animal, the “summed intensity” and PA were averaged among all ROIs. For each cohort, the mean “summed intensity” and mean PA were averaged among all animals.

### Pulse oximetry measurement of systemic oxygen saturation (SpO2)

The influence of LPS-induced inflammation on systemic arterial blood oxygen saturation was monitored in a separate group of APPswe/PS1dE9 and wildtype mice (*n*=3 each, at 6 months). The same LPS-administration protocol was applied (0.03 mg/kg LPS, injected intraperitoneally daily for 14 days). On each day, SpO2 and heart rate was measured on the mouse thigh using a commercial oximetry device (MouseOX Plus, Starr Life Science). Hair was removed to reduce measurement noise. During these measurements, the mice were placed under light anesthesia (1∼1.5% isoflurane) and kept warm with a heating pad. SpO2 were recorded for 20 s to 1 min. The time-averaged readout was used to represent systemic oxygenation on the measured day.

### Statistical analysis

All statistical analyses were performed using the Statistics toolbox from MATLAB. Two-tailed, independent student’s t-test was used to identify significant differences between AD and WT groups. Lillie’s test was used to evaluate the normality assumption. Bartlett’s test was used to check the equal variance assumption. Unequal variance was corrected with Satterthwaite’s approximation included in the “ttest2” function in MATLAB. Statistical significance level was set at *α*=0.05. One-way ANOVA with Tukey’s ‘hsd’ test was used for checking the difference between the cortical layers within each cohort. One-way repeated measures ANOVA with Bonferroni correction was used to analyze the effect of LPS-induced inflammation (66). Sphericity was checked using Mauchly’s test. Violation of sphericity was estimated by epsilon. Statistical significance level of ANOVA was set at 0.05. \**P*<0.05, \*\**P* <0.01, *** *P* <0.001. Sample size was determined using power analysis, assuming a 30% decrease of the measured physiological parameters (e.g., pO2) after induced inflammation, and an effect size of 0.8. *α* = 0.05.

## Results

We determined whether APPswe/PS1dE9 mice show differences in cortical oxygenation, microvascular hemodynamics, and neuroinflammation are evident at the early stages of AD progression using in vivo two-photon microscopy (TPM). We then explored how a 14-day LPS-induced neuroinflammatory stimulus alters neurovascular and metabolic regulation in AD progression.

### Arteriolar oxygen extraction in AD mice under baseline conditions

Prior to an LPS-induced inflammation (aka “baseline” conditions), we measured pO2 in diving arterioles and ascending venules from layers I to V in cortices of awake AD and WT mice at age 8 months (Fig. 2A). Our results showed that AD mice brains have slightly higher arteriolar and venular pO2 and SO2 compared to WT mice in all cortical layers (Fig. 2B). Statistically significant difference in pO2 was observed in arterioles at cortical layer IV, where the average pO2 in WT and AD is 69.33 ± 4.95 mmHg and 80.79 ± 1.65 mmHg, respectively. In cortical layer IV of AD mice brain, the comparable arteriolar SO2 but higher venular SO2 yielded significantly reduced depth-dependent oxygen extraction (DOE) in AD mice (Fig. 2B, *P*=0.0092 in layer IV). Numerous studies indicate that maximal oxygen demand occurs in cortical layer IV, which features the highest density of mitochondria as well as cytochrome oxidase.(67,68) Our results of reduced oxygen extraction in layer IV are indicative of diminished oxygen consumption by layer IV neurons in the presence of Amyloid β. Consistent with previous studies in awake 3–5-month-old female C57BL/6 mice (44,69), our measurements in WT mice showed lower arteriolar and venular pO2 at deeper cortical layers, which yields a higher oxygen extraction (DOE) in deep cortical layer IV (Fig. 2B).

**Fig. 2.**
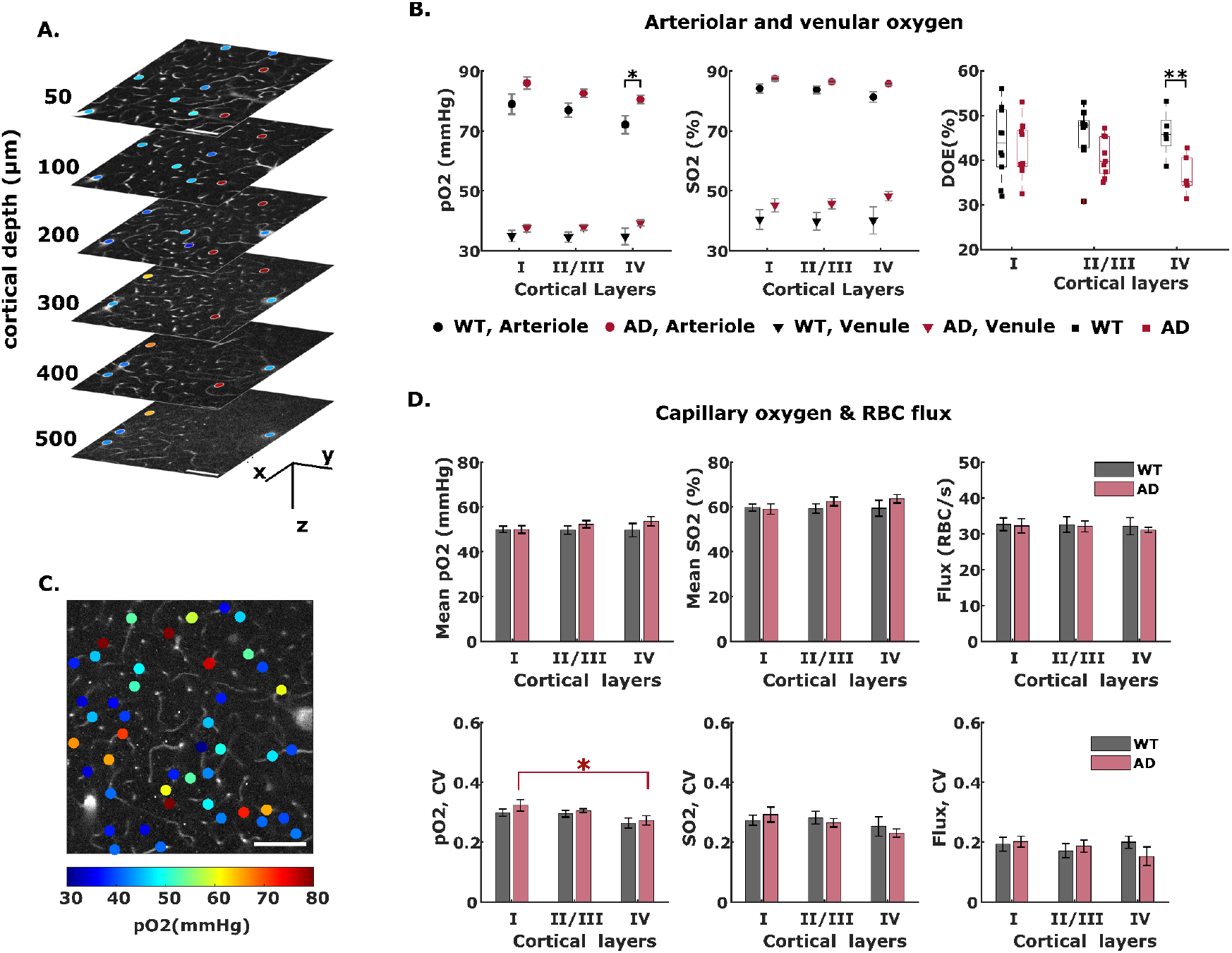
Baseline (pre LPS administration) measurement of cortical intravascular oxygenation and capillary flux in AD and WT mice. **(A)** 2D survey scan images of pO2 measurements in diving vessels at 50 µm to 500 µm below the pial surface (z=0 µm). scale bar = 100 µm. Colored dots are pO2-coded measurement points. **(B)** Cortical-layer dependent arteriole and venule pO2 and SO2, and cortical depth-dependent oxygen extraction (DOE) in AD and WT mice. Black asterisk symbol reveals the difference between the two cohorts of animals in cortical layer IV. **(C)** An example survey scan image of capillary pO2 measurement at cortical depth z=200 µm. Colored dots are pO2-coded measurement points. Colorbar is shared by **(A)** and (**C)**. **(D)** capillary oxygen and RBC flux. Upper row: cortical-layer-dependent capillary mean-pO2, SO2, and red blood cell (RBC) flux in AD and WT mice. Lower row: coefficient of variation (CV) of capillary pO2, SO2 and RBC flux of AD and WT mice. Data shown as mean ± standard error

### Baseline capillary pO2 and RBC flux are similar in AD and WT mice

Capillary pO2 and RBC flux were measured under baseline conditions at multiple cortical depths in AD and WT mice. Similar to the findings in arteriole and venule compartments, capillary pO2 and SO2 were slightly higher in AD mice compared to WT, without reaching statistical significance (Fig. 2D). Although our capillary pO2 observations in WT mice brains are slightly higher compared to previously reported values in C57BL/6J mice (48.59 ± 0. 31 mmHg vs 45.6 ± 1.4 in (44)), the differences are likely due to the different mouse strain and age. We calculated the coefficient of variation (CV) to evaluate the heterogeneity of capillary oxygen distribution (Fig. 2D). Consistent with previous findings by Li *et al* (44), we observed progressively lower oxygenation heterogeneity (indicated by lower CV) at deeper cortical layers in both WT and AD groups (Fig. 2D), with no appreciable difference between the two cohorts. Prior theoretical studies indicate that lower heterogeneity can promote oxygen extraction and maintain adequate oxygen supply to tissue (70). The decreased pO2 heterogeneity along cortical layers replicated previous results of more homogeneous oxygenation at deeper cortical layers in awake healthy mice (44).

We observed no significant differences between AD and WT mice in capillary RBC flux in any cortical layers (Fig. 2D), which agrees well with previous observations in a 6-month-old AD mouse model (71,72). RBC flux heterogeneity was also found to be similar between AD and WT mice (Fig. 2D). While decreased capillary flow and lack of capillary flow homogenization during functional activation has been observed in 18-month-old APPswe/PS1dE9 transgenic mouse (73), our results suggest that microcirculation is not profoundly altered at earlier ages.

### Neuroinflammation increases oxygen extraction in AD mice

LPS was applied daily for 14 days to assess how systemic inflammation affects cerebral oxygenation and hemodynamics in AD and wildtype mice. Fig. 3 illustrates how arterial and venous SO2 changes in diving arterioles and ascending venules over the course of the inflammatory threat. The same vessels were measured before, during and after induced inflammation (Fig. 3A). We averaged the measurements over animals to obtain the mean pO2 & SO2 in the two cohorts. Intriguingly, while systemic inflammation reduced microvascular oxygen levels in both cohorts, the effects were more pronounced and occurred earlier in AD mice (Fig. 3A-C, Additional file 1: Supplementary Fig. 1B-C). Within 7 days, LPS-induced inflammation substantially decreased arteriole and venule SO2 at all cortical layers in AD mice. By contrast, only mild, nonsignificant reductions were observed in WT mice at 7 days. In both cohorts, the corresponding changes to SO2 were most pronounced in ascending venules (Fig. 3B-C). The inflammatory threat induced substantial increases in computed depth-dependent oxygen extraction (DOE= SaO2 – SvO2) for AD mice in all cortical layers at 7 days of LPS administration (Fig. 3D). Conversely, in WT mice, significant increases in DOE were observed only in layer I at 7 days (Fig. 3D). At 14 days of LPS injection, both cohorts exhibited slight recovery in vascular oxygenation and oxygen extraction.

**Fig. 3.**
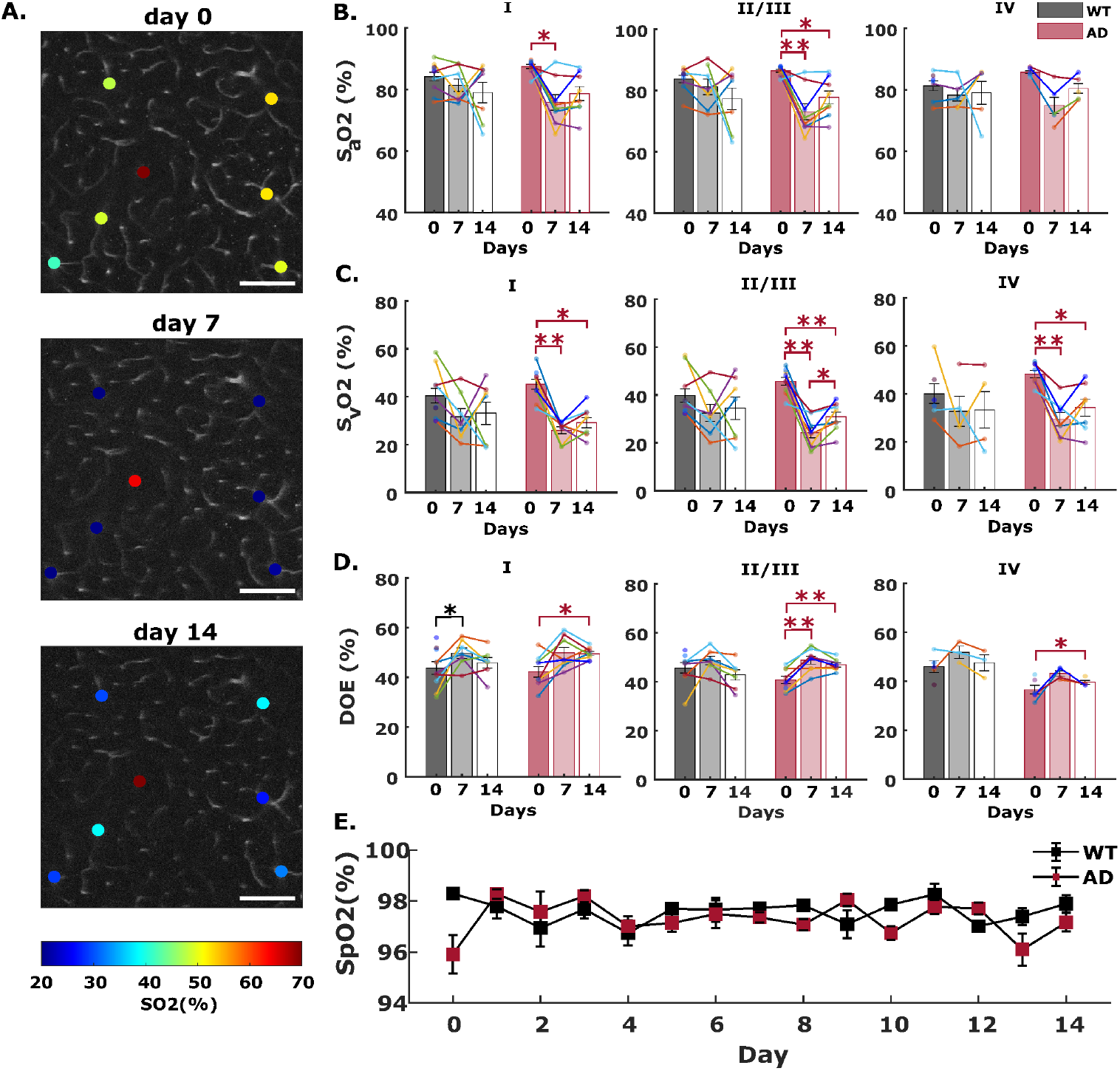
Inflammation-induced arteriole and venule oxygen reductions in WT and AD mice brain. **(A)** Example survey scan images overlaid with SO2 values in measured penetrating vessels at z=200 µm taken on day 0 (before LPS injection), day 7 and day 14 (with continuous LPS injection) inside an AD mouse brain. Scale bar = 100um. **(B-C)** Arteriole and venule SO2 measured on day 0, and day 7 and day 14 in WT and AD mice at cortical layer I, II/III, and IV. Each bar represents mean ± standard error over all measured mice in each cohort. Connected scatterplot shows the SO2 of each individual mouse. **(D)** Depth-dependent oxygen extraction (DOE) of WT and AD mice on day 0, day 7 and day 14. **(E)** Systemic oxygen saturation measurements during a neuroinflammatory stimulus, induced by 14-days of daily LPS administration (*n*=3 in each cohort). Steady SpO2 values impugn the prospect of LPS-induced impairments to pulmonary blood oxygenation

### LPS-induced inflammation does not impair pulmonary gas exchange

In AD mice, the significant oxygen reductions in O2-supplying arterioles at all cortical layers were unanticipated (Fig. 3B, and Additional file 1: Supplementary Fig. 1B). Prior studies in awake healthy mice indicate that only modest amounts of oxygen are delivered from cortical arterioles (44,62,65). Consequently, a reduction in this upstream vascular compartment warranted additional investigation. To determine whether the diminished oxygen in cortical arterioles results from impaired pulmonary gas exchange, we monitored the systemic arterial oxygenation (SpO2) in a separate group of AD and WT mice during 14 days of induced systemic inflammation. Insufficient oxygen uptake in the lung compartment would yield global reductions in oxygen supply and therefore cause substantially lower SpO2. Our daily measurements from the mouse thigh revealed no appreciable changes of SpO2 in either mouse cohort during the course of a 14-day inflammatory threat (Fig. 3E), which belies the possibility of diminished pulmonary gas exchange induced by systemic inflammation.

### LPS-induced inflammation decreases capillary pO2 but not capillary RBC flux

Capillary pO2 measurements were performed in the same fields of view on days 0, 7 and 14 of LPS-induced inflammation (Additional file 1: Supplementary Fig. 2), and corresponding SO2 values were calculated using the Hill equation. Fig. 4 displays how LPS-induced inflammation alters capillary SO2 and their associated coefficients of variation (CV) at different cortical layers. After 7 days, notable capillary SO2 decreases were observed in all cortical layers of WT brain (Fig. 4B-C). In AD mice, the inflammatory threat yielded more drastic reductions (Fig. 4E). In both AD and WT cohorts, the most pronounced oxygen reductions were observed in cortical layer IV (Fig. 4E, ΔSO2= −13.22 ± 3.36 % in WT and ΔSO2= −23.17 ± 3.28 % in AD). Capillary oxygenation became substantially more heterogenous in both cohorts under inflammatory threat, as indicated by the increased CVs of capillary SO2 (Fig. 4D). Prior modeling studies indicate that greater dispersity of capillary transit time markedly limits total O2 extraction from blood (44,70,75). Guided by their findings, our pO2 and SO2 heterogeneity observations strongly indicate that systemic inflammation substantially disrupts the delicate balance between oxygen supply and demand in cortical brain tissue.

**Fig. 4.**
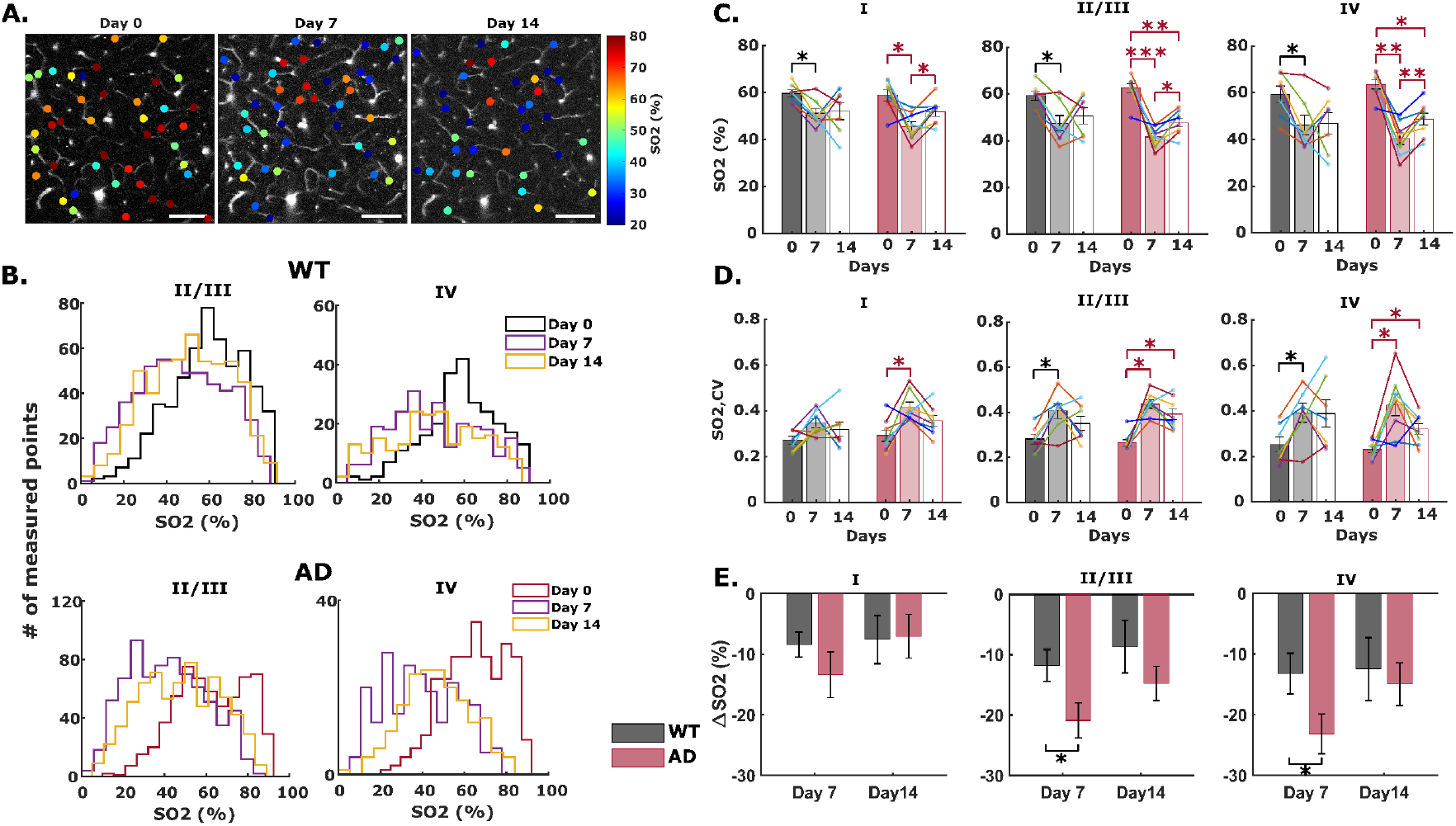
Inflammation-induced cerebral capillary oxygen reduction in WT and AD mice brain. **(A)** Example survey scan images overlaid with SO2 values in measured capillary segments at z=200 µm taken on day 0 (before LPS injection), day 7 and day 14 (with continuous LPS injection) inside an AD mouse brain. Scale bar = 100um. **(B)** Histogram showing layer-specific SO2 distributions in all measured capillary points from all animal samples in both cohorts. **(C-D)** Capillary SO2 and the corresponding coefficient of variation (CV) at baseline (day 0), and with LPS-induced inflammation (day 7 and 14) in WT and AD mice brain in cortical layer I, II/III, and IV. Each bar represents mean ± standard error over all measured mice in each cohort. Connected scatterplot shows the SO2 of each individual mouse. **(E)** Absolute changes in capillary SO2 in WT and AD mice with 7 and 14 days of LPS injection. *ΔSO2* is calculated by subtracting the SO2 on day 0 from the SO2 on day 7 or 14.

Interestingly, after 14 days of LPS injection, cerebral capillary oxygenation and oxygen heterogeneity demonstrated a slight tendency to recover in both mice cohorts (Fig. 4C-D, and Additional file 1: Supplementary Fig. 2). This observation motivated additional investigations into the duration of inflammatory disruptions. In a subset of the remaining animals (*n* = 2 AD, *n*= 2 WT), arteriolar, venular, and capillary oxygen measurements recovered to the baseline level after 70 days (∼10 weeks) since the final LPS injection (Additional file 1: Supplementary Fig. 3). Additional investigations will explore the extent of reversibility of inflammation’s metabolic disruptions.

In contrast to its effects on intravascular oxygenation, systemic inflammation did not significantly affect layer-dependent capillary RBC flux in AD or WT mice (Fig. 5). In WT mice, we observed mild, but not significant reductions in RBC flux within 7 days in layers II – V. The changes were even less pronounced in AD mice (Fig. 5A-B). On average, the coefficients of variation for RBC flux did not change appreciably after systemic inflammation (Fig. 5C), suggesting that spatial blood flow heterogeneity remains largely unaffected by an inflammatory threat in 8-month-old mice.

**Fig. 5.**
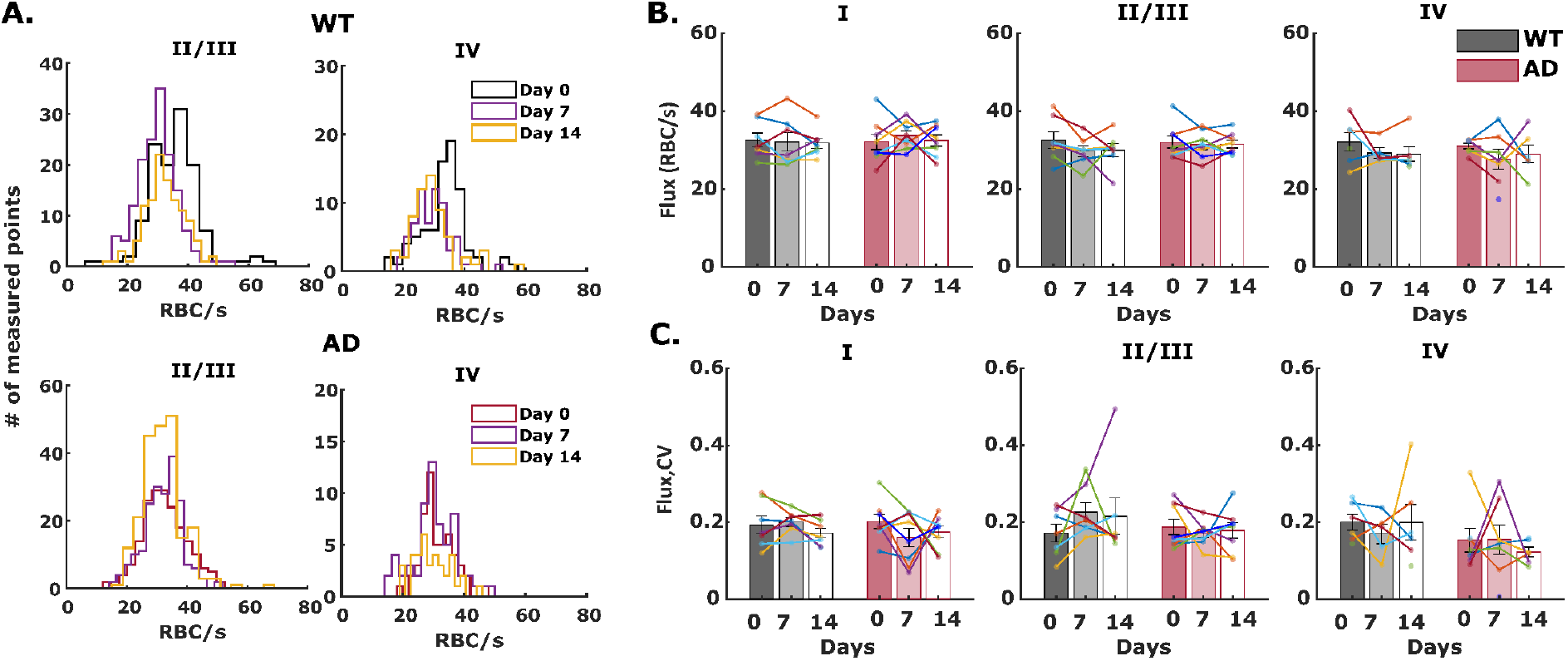
Cerebral capillary red blood cell flux (RBC flux) at day 0 (no LPS injection), day 7 and day 14 (with LPS injection) in WT and AD brain. **(A)** Histograms showing RBC flux in all measured capillary points in cortical layers II to IV. **(B-C)** Mean RBC flux and coefficient of variation over all measured mice in both cohorts over the course of LPS-induced inflammation. Each bar represents mean ± standard error over all measured mice in each cohort. Connected scatterplot shows the RBC flux and the CV of each individual mouse

### LPS induces inflammatory secretions in microglia and astrocytes

Glial fibrillary acidic protein (GFAP) and ionized calcium-binding adaptor molecule (Iba-1) are biomarkers of microglia and astrocytes, respectively, and are upregulated in activated microglia and astrocytes during neuroinflammation (76,77). We quantified the activation of microglia and astrocytes through immunofluorescence staining of Iba1 and GFAP in AD and WT mice after 14 days of daily LPS administration. Somatosensory cortex in both hemispheres were imaged with confocal microscopy (Additional file 1: Supplementary Fig. 4A). Additional file 1: Supplementary Fig. 4b shows representative maximum intensity projections (MIP) of 40µm-thick z-stack images. Prior to inflammation, there were no significant differences in the expression of Iba1 and GFAP between WT and AD mice (Additional file 1: Supplementary Fig. 4C-D), consistent with previous immunological studies showing modest, nonsignificant inflammation in 6-month-old APP/PS1dE9 mice cortex (78,79). Elevated expression of Iba1 and GFAP was observed in both cohorts after 14 days of LPS-induced inflammation, as indicated by increased percentage area and total fluorescence intensity of the Iba1 and GFAP (Additional file 1: Supplementary Fig. 4C-D). Compared to WT, AD mice showed more pronounced increases in Iba1 and GFAP, indicating a pronounced neuroinflammatory response.

## Discussion

Neuroinflammation reportedly has both therapeutic and harmful effects during AD progression (1,40). The conflicting influences of neuroinflammation have motivated several studies exploring the effects of modulating neuroimmunity (41). The impact of a pronounced inflammatory threat on cerebral energetics and microvascular hemodynamics warrants much further investigation to guide effective diagnostic and therapeutic developments. To our knowledge, we provide the first quantitative measurements of microvascular oxygen pressure, oxygen extraction, and RBC flux in the APP/PS1dE9 mouse cortex before, during, and after a pro-inflammatory stimulus.

Under baseline conditions (in the absence of inflammatory stimulus), our results in WT mice indicate that depth-dependent oxygen extraction (DOE) increases along cortical layers, consistent with previous 2P-PLIM measurements in awake C57 mice (Fig. 2) (44,69). AD mice have slightly higher pO2 and oxygen saturation in arterioles, capillaries, and venules at all cortical layers, corresponding to progressively lower DOE at deeper cortical layers. Conversely, no significant differences in capillary RBC flux were found in either AD or wildtype cohorts at any cortical layer. Taken together, these observations suggest that early amyloid accumulation induces reductions in cortical oxygen extraction without yet substantially altering vascular oxygen supply in 8-month-old APPswe/PS1dE9 mice. Our results suggest that cerebral metabolic rate of oxygen (CMRO2 = CBF * OEF) would be reduced in AD mice, primarily due to reductions of the brain’s metabolic demand.

Our baseline observations in AD mice agree well with human MRI-studies in patients with amnestic mild cognitive impairment (MCI, regarded as an early stage of AD) (80). Global cerebral blood flow was found to be similar in MCI patients relative to age-matched control patients, while OEF was lower in MCI patients by ∼10%. Consequently, with no notable changes to blood supply, the authors attributed the ∼12.5% lower global CMRO2 in MCI patients to a reduction in metabolic demand by the brain tissue. Also consistent with our results, the same authors reported that OEF was reduced in later-stage AD patients, and the magnitude of OEF reduction correlated well with severity of cognitive impairment (81).

In our results, the significantly lower DOE in cortical layer IV in AD mice indicates lower demand of oxygen by layer IV neurons, which is likely the result of reduced neural activity. This correlates well with observations of reduced mitochondrial function in AD mice at age 7-10 months (82) and reduced EEG rhythms observed in MCI patients (80). The lower DOE in our 8-9-month-old AD mice are likely a consequence of Aβ-induced impairments to mitochondrial function and diminished neuronal activity (82–84). Contrast-enhanced MRI measurements in APPswe/PS1dE9 mice showed no changes in relative cerebral blood volume (rCBV) at 8-month-old mice but significant reduction at 15-month-old mice (85). We expect that APPswe mice would likely also experience pronounced impairments to CBF at more advanced ages that exhibit more severe amyloid pathology. Future investigations will explore how oxygen extraction and other metabolic markers vary at more advanced ages in AD mice, as well as age-dependent susceptibility to systemic inflammatory threats.

While our observations indicate that baseline oxygen extraction in AD mice is lower than WT mice, an inflammatory insult provoked increased extraction of oxygen in both animal groups. More pronounced changes to SO2 and DOE were consistently observed in AD mice (Fig. 3 and 4). Conversely, we found no significant inflammation-induced changes to RBC flux in capillaries in cortical layers I-IV (Fig. 5). Our results imply that, at the onset of old age in mice (7-9 months), systemic inflammation substantially increases oxygen demand by the brain tissue without affecting vascular supply of metabolites.

We posit that the increased oxygen demand is the net result of multiple pathological processes triggered by an inflammatory insult. Increased metabolic demand by hyperactive neurons most likely accounts for the majority of increased O2 consumption during inflammation (86). However, in addition to increased energetic demand, additional amounts of oxygen are also extracted from blood to facilitate multifaceted actions of immune cells. Numerous studies have demonstrated that LPS administration yields dramatic increases in spontaneous calcium spiking and excitatory post-synaptic currents from neurons in the cortex, hippocampus, and several other regions of the brain, while also reducing the activity of inhibitory GABAergic interneurons (55,86–90). Because restoration and maintenance of ion gradients accounts for the bulk of ATP consumption in the brain, and energy consumption increases markedly with increased action potential frequency, inflammation-induced neuronal hyperactivity across different brain regions constitutes a massive additional energetic burden (91). In addition to its effect on neuronal hyperactivity, though, LPS-induced neuroinflammation triggers a host of actions by immune cells with varied effects on cerebral oxygen demand. Upon activating to their pro-inflammatory phenotype to combat pathological threats, microglia and astrocytes avidly produce inflammatory cytokines, increase phagocytic activity, and undergo morphological changes to more ameboid profiles. To facilitate cell proliferation and rapid cytokine production, activated microglia reportedly shift their metabolism to rely more on glycolysis, while also experiencing increased mitochondrial fission (30,33,52). While the former adaptation reduces microglial oxygen dependence for metabolism, investigators observed that LPS exposure actually increases oxygen consumption rate (OCR) in microglia, particularly OCR associated with mitochondrial proton leak (92). Despite their purported shift to glycolytic metabolism, our observations support prior studies, which demonstrate that activated microglia consume higher amounts of oxygen to mediate ROS production to neutralize pathogenic threats. Microglia show similar metabolic behavior to peripheral immune cells such as monocytes, leukocytes, and neutrophils, which also contribute to the neuroinflammation response and can increase ROS generation by 50-fold during high immune activity (28,30,93–95).

Overall, our data demonstrate that the increased oxygen demand from hyperactive neurons and pro-inflammatory immune cells is substantial enough to extract oxygen from the arteriolar compartment. In contrast to the classical Krogh model of exclusive oxygen extraction from the capillary compartment, significant oxygen extraction from arterioles has been observed previously in healthy mice under isoflurane anesthesia. In the absence of anesthesia, though, similar measurements in awake mice agree better with the Krogh model (44,65,69,46). Our observations indicate that pronounced neuroinflammation substantially, but reversibly, shifts the balance of metabolite supply and demand in the brain.

Compared to wildtype mice, the more aggravated response in AD mice can be attributed to a deteriorative cycle of cytokine release and impaired mitochondrial function, initiated and promoted by continued accumulation of cytotoxic amyloid β. A prior study showed that APP/PS1dE9 mice at age 9-12 months secrete significantly higher TNF and IL-1β than wildtype mice in response to long-term systemic LPS injection (96). Similarly, our IHC results indicate that LPS triggers greater release of IBA-1 and GFAP from AD mice (Additional file 1: Supplementary Fig. 4), supporting the assertion that AD mice favor proinflammatory phenotype upon innate immune stimulus. Prior reports in both humans and mice also demonstrate links between elevated levels of proinflammatory cytokines (e.g., TNF-α) and increased both tau and cytotoxic amyloid pathology (97–99). In essence, a positive feedback loop exists between amyloid pathology and neuroinflammation that further exacerbates the brain’s pathological disruptions. Energy metabolism and oxygen demand contribute to this destructive feedback loop through inflammatory signaling’s marked impairments to mitochondrial function. Cytokines such as TNFα and IL-1β as well as ROS disrupt mitochondrial enzyme activity involved in the TCA cycle, inhibits oxidative phosphorylation (OXPHOS), and reduces ATP production (20). LPS reportedly induces both short- and long-term reductions on OXPHOS by downregulating activity of complexes I, II, and IV. While the exact mechanisms are not precisely understood, one proposed model hypothesizes that the LPS-induced torrent of cytokines such as TNFα activate a downstream tyrosine kinase, leading to the phosphorylation of complex IV and strong inhibition of all electron transport chain activity. Ultimately, impairments to the TCA cycle and electron transport chain significantly reduces mitochondrial membrane potential and efficiency of oxidative phosphorylation (100). Leaked components from dysfunctional mitochondria could in turn further activate innate inflammatory response and impair Aβ clearance (101). A vicious cycle ensues between Aβ, inflammation, mitochondrial dysfunction, and oxidative stress in AD mice brain that lower the efficiency of oxygen usage and ATP synthesis. As a result, tissue oxygen extraction is greatly increased to sustain the heightened energy needs and inflammatory response in the AD brain (Fig. 3).

In both cohorts, induced inflammation did not significantly affect cerebral capillary RBC flux (Fig. 5). Our results are consistent with previous reports of no change of cerebral blood flow after LPS-induced acute inflammation (102,103). Fruekilde *et al* reported increased resting-state capillary stalling under LPS-induced acute inflammation due to leukocyte plugging (104).

## Conclusions

This study demonstrates that induced systemic inflammation reduces cerebral oxygen consumption without disturbing microvascular supply of oxygen in a mouse model of Alzheimer’s disease at the age of 8-9 months. Our findings substantiate purported links between neuroinflammation and energy metabolism in preclinical AD pathology. The results will help advance the understanding of the critical role inflammation plays in Alzheimer’s disease and the utility of oxygen extraction fraction (OEF) in AD diagnosis. Our preclinical investigation provides insight to better interpret observations from translational neuroimaging using modalities such as functional magnetic resonance imaging (fMRI) and positron emission tomography (PET). And the interaction between AD brain immune response and cerebral oxygen metabolism should be taken into account when developing AD treatment targeting neuroinflammation. One limitation of our study is that we did not label Aβ. Further investigations will explore longitudinal labeling of Aβ during systemic inflammation to correlate the oxygenation changes with amyloid pathology. Additionally, though we observed no significant differences in baseline capillary RBC flux and spatial heterogeneity, our results do not comprehensively refute prospective impairments to CBF and microvascular hemodynamics at early stages of AD. Our observations are consistent with a prior study of capillary RBC flux in the barrel cortex of 6-month APP/PS1dE9 mice (71). Hypoperfusion is widely recognized as an early pathological indicator of AD, but conflicting studies with different animal models, different ages, and different measurement techniques confound a detailed understanding of the precise timing, severity, and spatial heterogeneity of CBF impairments during the preclinical stage of AD (12,71,105,106). Some studies showed no obvious changes in CBF and capillary flow in young mice but severe reduction of flow and disturbed flow homogenization in aged transgenic mice (73). Recent advances in optical coherence tomography (OCT) will help delineate the precise trajectory of hemodynamic alterations during AD by enabling near-lifespan tracking of microvascular morphology and function (107). Lastly, estimating cerebral metabolic of oxygen (CMRO2) in response to inflammation using Krogh-Erlang model (108) or the recent proposed ODACITI model (109) could further validate our findings of increased oxygen extraction in inflamed AD brain tissue.

## List of abbreviations

2P-PLIM: Two-photon phosphorescence lifetime imaging microscopy
AD: Alzheimer’s disease
APP: Amyloid precursor protein
ATP: Adenosine triphosphate
Aβ: amyloid-beta
CBF: Cerebral blood flow
CMRO2: Cerebral metabolic rate of oxygen
CV: Coefficient of variation
DOE: Depth-dependent oxygen extraction
fMRI: Functional magnetic resonance imaging
GFAP: Glial fibrillary acidic protein
IBA-1: Ionized calcium-binding adaptor molecule 1
LPS: Lipopolysaccharide
NGVU: Neuro-glio-vascular unit
OCT: Optical coherence tomography
PET: Positron emission tomography
pO2: Partial pressure of oxygen
PS1dE9: Presenilin 1 gene
RBC: Red blood cell
ROS: Reactive oxidative species
SO2: Oxygen saturation
TCA: Tricarboxylic acid cycle

## Declarations

### Ethics approval and consent to participate

All experiments were performed in accordance with ARRIVE guidelines for animal care, under a protocol approved by the Northeastern University Institutional Animal Care and Use Committee.

### Consent for publication

Not applicable

### Availability of data and materials

Data generated and analyzed in the study are available upon request.

### Authors’ contributions

CL and MAY conceptualized and designed the study. CL performed animal surgery, collected and analyzed the 2P-PLIM data. ST and AB conducted confocal imaging and analyzed the data. CL, ACR and MA developed data analysis protocol. CL and MAY wrote the first draft of the manuscript. All authors commented on and approved the final manuscript.

## Acknowledgements

We thank Dr. Sava Sakadžić (Massachusetts General Hospital, Harvard Medical School) for the discussion on pO2 calculation and Dr. Anna Devor (Boston University) for the productive discussions on our results. We thank Dr. Qi Pan (Massachusetts General Hospital, Harvard Medical School) and Dr. John Giblin (Boston University) for the approach to capillary flux calculation. We thank Dr. Buyin Fu (Massachusetts General Hospital, Harvard Medical School) and Dr. Kivilcim Kilic (Boston University) for guidance with animal preparation. We thank Dana-Farber/Harvard Cancer Center in Boston, MA, for the use of the Specialized Histopathology Core, which provided brain slices immunofluorescence staining service. Dana-Farber/Harvard Cancer Center is supported in part by an NCI Cancer Center Support Grant # NIH 5 P30 CA06516.

