## Additional File 1 for "Neuroinflammation increases oxygen extraction in a mouse model of Alzheimer’s disease"

**Supplementary Table 1. Number of arterioles, capillaries, venules and number of animals (n) measured for pO2.**

| Cohort | | WT | | | AD | | |
|---|---|---|---|---|---|---|---|
| Measurement | Vessel type | Layer I | Layer II/III | Layer IV | Layer I | Layer II/III | Layer IV |
| Baseline | arterioles | 22 (10) | 21 (9) | 15 (7) | 18 (9) | 18 (9) | 8 (6) |
| | venules | 43 (10) | 43 (10) | 29 (6) | 44 (9) | 44 (10) | 28 (8) |
| | capillaries | 461(7) | 549(7) | 252(7) | 497 (8) | 606 (8) | 269 (8) |
| Day 7 | arterioles | 13 (7) | 10 (7) | 6 (5) | 16 (8) | 15 (8) | 6 (5) |
| | venules | 27 (7) | 27 (7) | 8 (4) | 37 (8) | 35 (8) | 15 (7) |
| | capillaries | 512(7) | 548(7) | 247(7) | 701(8) | 735 (8) | 223 (8) |
| Day 14 | arterioles | 16 (7) | 14 (7) | 6 (5) | 16 (8) | 14 (8) | 5 (5) |
| | venules | 30 (7) | 29 (7) | 8 (4) | 37 (8) | 33 (8) | 17 (7) |
| | capillaries | 537(7) | 551(7) | 218(7) | 609 (8) | 695 (8) | 179 (8) |

**Supplementary Table 2. Number of capillaries and animals (n) measured for RBC flux.**

| Cohort | WT | | | AD | | |
|---|---|---|---|---|---|---|
| Measurement | Layer I | Layer II/III | Layer IV | Layer I | Layer II/III | Layer IV |
| Baseline | 104 (7) | 122 (7) | 79 (6) | 162 (8) | 191 (8) | 42 (6) |
| Day 7 | 146 (7) | 156 (7) | 66 (6) | 163 (8) | 200 (8) | 60 (6) |
| Day 14 | 147 (7) | 116 (7) | 60 (6) | 145 (8) | 219 (8) | 33 (6) |

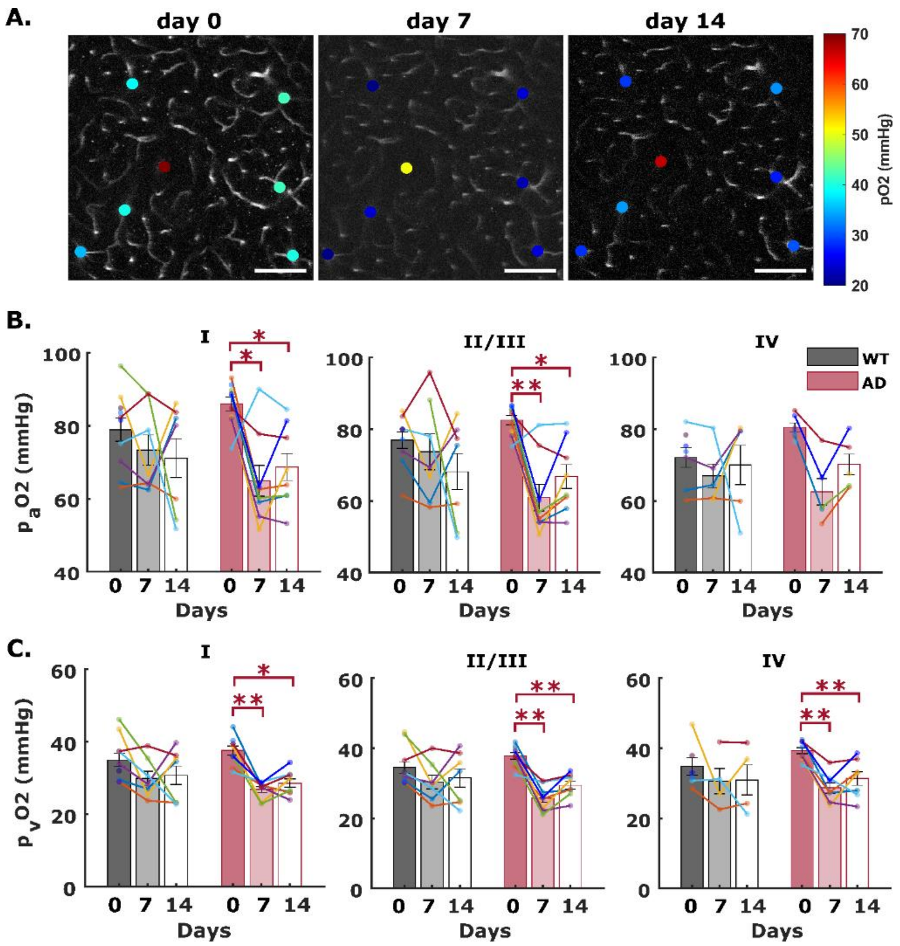


**Supplementary Fig. 1 Inflammation-induced arteriole and venule pO2 reductions in WT and AD mice brain. (A)** Example survey scan images overlaid with pO2 values in measured penetrating vessels at z=200 µm taken on day 0 (before LPS injection), day 7 and day 14 (with continuous LPS injection). Scale bar = 100um. **(B-C)** Arteriole and venule pO2 measured on day 0, and day 7 and day 14 in WT and AD mice at cortical layer I, II/III, and IV. Each bar represents mean ± sem over all measured mice in each cohort. Connected scatterplot shows the SO2 of each individual mouse

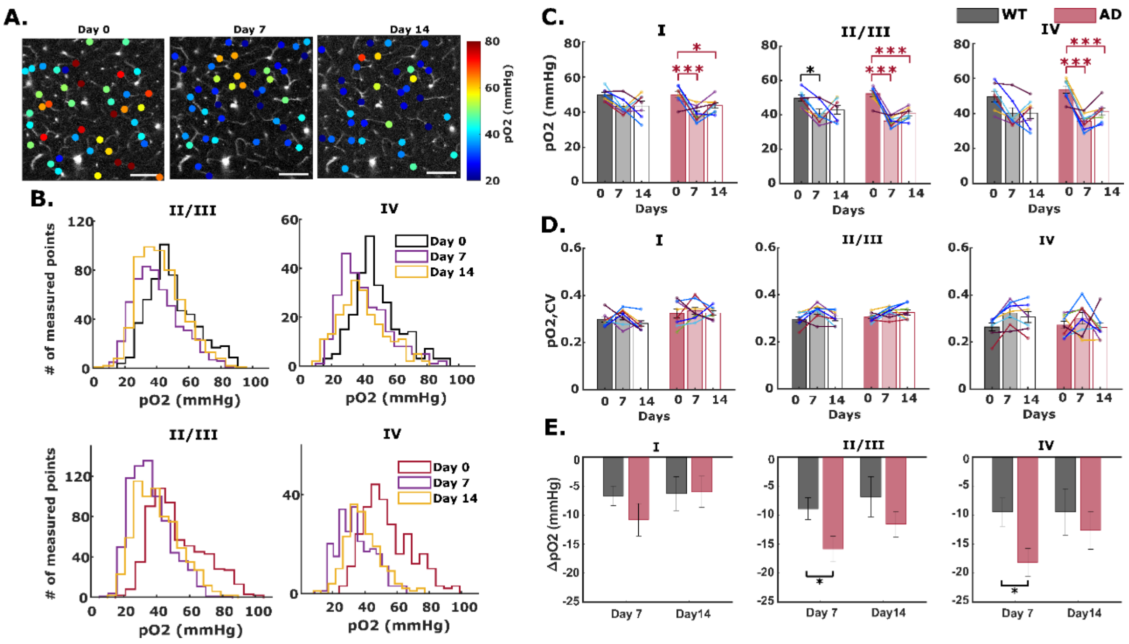


**Supplementary Fig. 2 Inflammation-induced cerebral capillary oxygen reduction in WT and AD mice brain.** **(A)** Example survey scan images overlaid with pO2 values in measured capillary segments at z=200 μm taken on day 0 (before LPS injection), day 7 and day 14 (with continuous LPS injection). Scale bar = 100um. **(B)** Histogram showing layer-specific pO2 distributions in all measured capillaries at cortical layer II to IV. **(C-D)** Capillary pO2 and the corresponding coefficient of variation (CV) at baseline (day 0), and with LPS-induced inflammation (day 7 and 14) in WT and AD mice brain at cortical layer I, II/III, and IV. Each bar represents mean ± standard error over all measured mice in each cohort. The connected scatterplot shows the SO2 of each individual mouse. **(E)** Absolute changes in capillary pO2 in WT and AD mice with 7 and 14 days of LPS injection

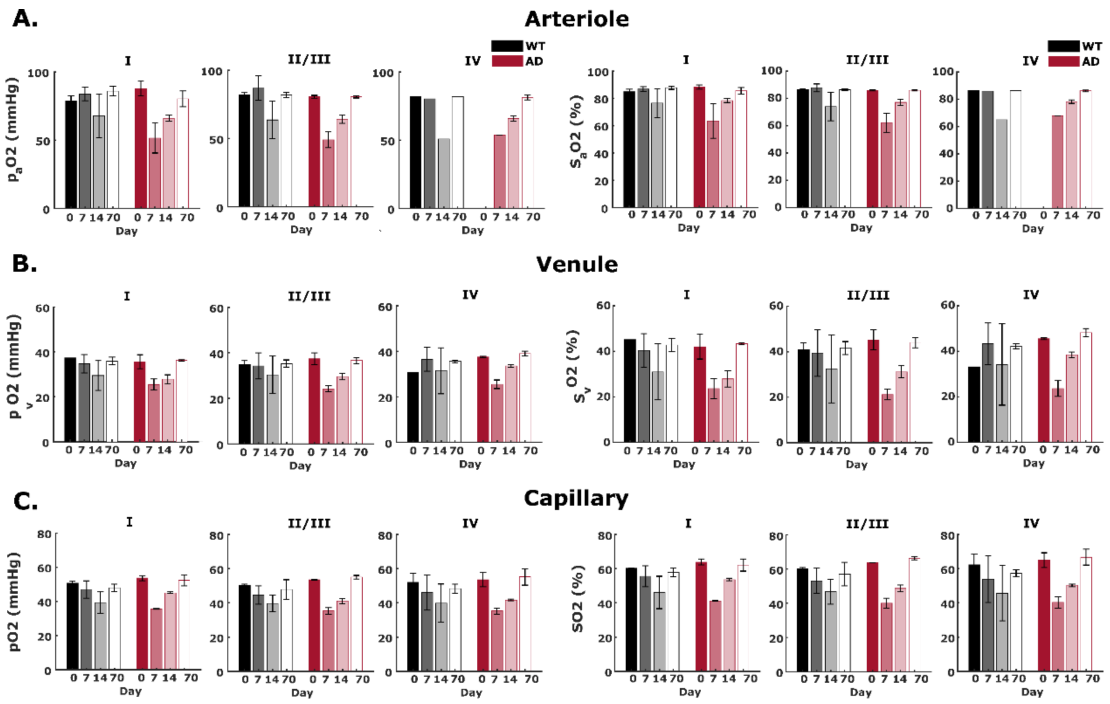


**Supplementary Fig. 3 Recovery of cerebral intravascular oxygenation in WT and AD mice (*n*=2 in each cohort).** pO2 and SO2 in arterioles **(A)**, venules **(B)**, and capillaries **(C)** were measured 70 days after the last LPS injection. pO2 and SO2 in all three vessel types in the measured mice were recovered to the same level as baseline (before LPS-induced inflammation). Data displayed as the mean pO2 (SO2) in cerebral arteriole, venules, and capillaries over two AD and two WT mice. Error bar represents the standard error

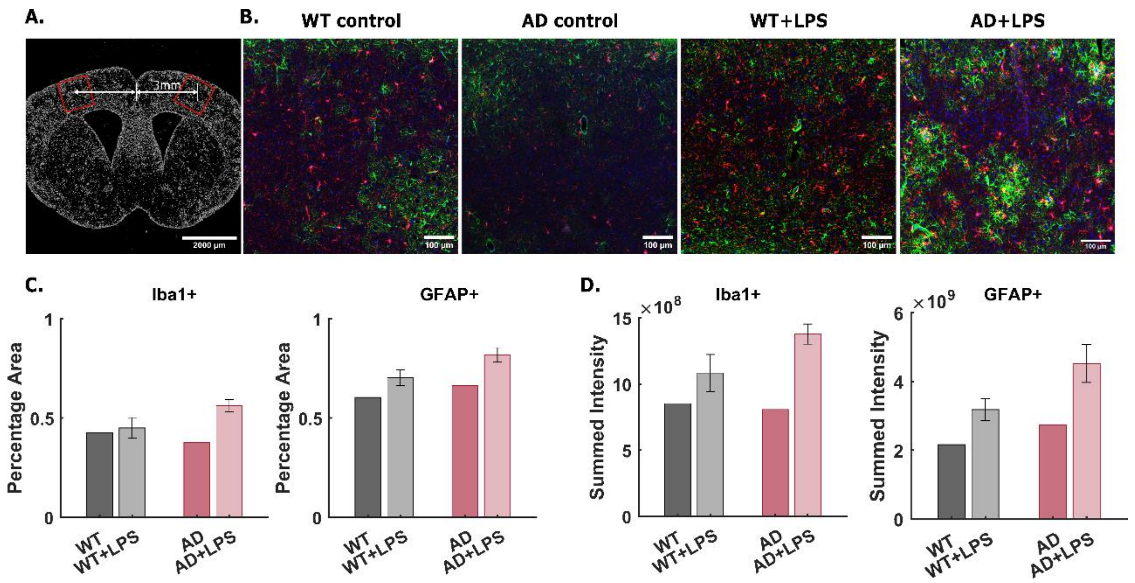


**Supplementary Fig. 4 (A) Example coronal brain sections with the analyzed FOV indicated in red rectangle boxes. Fluorescence signal is DAPI-labelled nuclei. (B)** Example maximum intensity projection (MIP) of confocal images of WT and AD mice. Control groups are mice that did not receive LPS injection. Green: astrocyte. Red: microglia. Blue: nuclei. Images displayed have the same intensity threshold. **(C)** Percentage area (%) of Iba1 positive microglia and GFAP positive astrocyte in the image field. **(D)** Summed fluorescence intensity showing the expression of Iba1 in microglia cells and GFAP in astrocytes. Note that all images are imaged with same settings of excitation power and PMT gain and are not saturated. Data shown as mean ± standard error. *n*=1 in WT and AD without LPS injection, *n*=3 in WT and AD with LPS injection
